# ENVELOPED VIRUS-LIKE PARTICLES (eVLPs) EXPRESSING MODIFIED FORMS OF ZIKA VIRUS PROTEINS E AND NS1 PROTECT MICE FROM ZIKA VIRUS INFECTION

**DOI:** 10.1101/666966

**Authors:** Anne-Catherine Fluckiger, Jasminka Bozic, Abebaw Diress, Barthelemy Ontsouka, Tanvir Ahmed, Amalia Ponce, Marc Kirchmeier, Francisco Diaz-Mitoma, Wayne Conlan, David E. Anderson, Catalina Soare

**Affiliations:** VBI Vaccines, Cambridge, 222 Third Street, Cambridge, 02142 Massachusetts, USA; National Research Council Canada, Department of Human Health Therapeutics, 100 Sussex Drive, Ottawa ON K1A0R6, Canada

## Abstract

While Zika virus (ZIKV) infection induces mild disease in the majority of cases, it has been identified as responsible for microcephaly and severe neurological disorders in recent 2015-2016 outbreaks in South America and the Caribbean. Since then, several prophylactic vaccine strategies have been studied. Here, we describe the development of a ZIKV candidate vaccine consisting of bivalent enveloped virus-like particles (eVLPs) expressing a modified form of E and truncated NS1 (EG/NS1) proteins. In EG/NS1, the E transmembrane/cytoplasmic tail has been replaced with those domains from the VSV G protein and a β-domain of NS1 was fused in-frame to Gag from Moloney murine leukemia virus (MLV). Immunization of BALB/C mice demonstrated that bivalent EG/NS1 and monovalent EG eVLPs induced comparable levels of antibody (Ab) titers but that EG/NS1 induced much higher neutralizing activity, comparable to naturally acquired anti-ZIKV immunity. In contrast, monovalent NS1 eVLPs did not induce a significant anti-NS1 Ab response but promoted strong T cell immunity that was also elicited with EG/NS1 eVLPs. ZIKV challenge studies in C57BL/6-IFNαR^−/−^ mice demonstrated that EG/NS1 eVLPs conferred 100% protection against clinical disease after ZIKV challenge compared to 80% protection after EG eVLP vaccination, with protection against challenge correlating with neutralizing antibody titers and overt signs of infection.

**Author Summary:** Zika virus has caused rapidly spreading epidemics with potentially severe neurological symptoms including microcephaly in new born babies. Rapid progress has been made with several candidate vaccines under clinical evaluation but no vaccine or treatment is yet available. In this context, we have produced and tested recombinant virus-like particles that incorporate one or two Zika virus proteins, E and NS1 that have been modified for optimal efficacy. Our immunogenicity studies in mice showed a synergistic effect of both proteins in the bivalent vaccine. NS1 induced a strong T cell response enhancing the neutralizing antibody production induced by the E protein. In challenge experiments, the bivalent vaccine protected 100% of mice from clinical signs of Zika virus infection. These products could be further used to explore Zika virus correlates of protection and evaluated as vaccine candidates.

## Introduction

Zika virus was first isolated in 1947 in a Rhesus sentinel monkey in Uganda, and was not initially regarded as a significant human pathogen. However, large outbreaks in the Pacific area including Yap Island in 2007; French Polynesia in 2013 to 2014) and the intense epidemic in The Caribbean and South America in 2016 (1,2), led the WHO to declare ZIKV a global public health emergency and the development of a vaccine a priority. Despite the end of the outbreak and the ongoing decline of the number of cases, the need for a licensed ZIKV vaccine remains critical for global public health. Indeed, the ZIKV infection is generally asymptomatic or induces a mild fever, but is also associated to severe neurological syndromes such as Guillain Barré Syndromes in adults, and microcephaly in infants. CDC recently established that about one of seven children born to mother with ZIKV infection during pregnancy had Zika-associated birth defect, a neurodevelopmental abnormality possibly associated with congenital Zika virus infection, or both (3). In contrast to other described arbovirus infections, ZIKV RNA can persist in human semen for a year or more, and cases of sexual transmission have been reported (4). Cultured human germ cells are permissive to ZIKV infection and can produce infectious virus (5). These latter features of ZIKV could have impacts on demographics (6). There are still no licensed vaccines or therapeutic measures against ZIKV infection.

More than forty ZIKV vaccine candidates have advanced through efficacy studies in mice and non-human primates (NHP), nine progressed in Phase I clinical trials and among them two entered Phase II. Early findings like current efforts suggested that the presence of high titers of neutralizing antibodies against the ZIKV envelope protein E correlate with protection (7–9). In addition, more recent studies have exposed an important role of CD4^+^ T-cell activation to develop and maintain an efficient protective response during ZIKA infection (10,11). In accordance with these findings, a T-cell biased adenovirus vector expressing prME has been shown to protect mice against ZIKV challenge without detectable levels of ZIKV specific Ab. Finally, ZIKV vaccine candidates based on expression of the non-structural protein NS1 have also shown protection of mice from ZIKV infection by inducing both humoral and cellular responses (12).

The prME polyprotein is expressed in immature ZIKV as trimeric spikes of prM:E heterodimers, with the pr peptide covering the fusion loop. In mature particles, cleavage of pr and rearrangement of the glycoproteins, lead to a smooth virion enveloped by trimers of E:M heterodimers organized in an herringbone like symmetry (13,14). ZIKV and dengue virus (DENV) share high degrees of homology not only in their amino-acid sequences, but also on the structure of the E protein at the surface of the virus, resulting in production of cross-reactive antibodies (15–19).

There is evidence of antibody-dependent enhancement (ADE) of infection that occurs in the presence of cross-reactive but weakly neutralizing Abs (nAb) (20). Given that ZIKV and DENV are endemic in similar regions, an important consideration in ZIKV vaccine development is generation of an efficacious vaccine which does not inadvertently enhance the risk for ADE associated with DENV infection. eVLPs enable repeating, array-like presentation of antigens which is a preferred means of activating B cells and eliciting high affinity antibodies (21). The potential to enrich for antibodies with neutralizing and not just binding activity may help avoid the potential for ADE. We describe here the development of ZIKV vaccine candidates, expressing a modified ZIKV E protein alone or in combination with NS1, and their relative abilities to induce neutralizing antibody responses in mice and protection from ZIKV challenge.

## Results

### Production and characterization of eVLPs expressing high densities of ZIKV E protein

We tested the feasibility of expressing ZIKV E protein on the surface of MLV Gag-based eVLPs. In previous studies, we and others have observed that fusion of the ectodomain of a viral glycoprotein with the transmembrane/cytoplasmic (TM/Cyt) domain from VSV-G promoted pseudotyping of MLV Gag pseusoparticles (24,25). We tested a similar strategy for ZIKV E and compared expression of EG to expression of full length prME in eVLPs.

HEK 293 cells were cotransfected with the MLV Gag plasmid and an expression plasmid coding either for the full-length ZIKV prME sequence or for a modified form of ZIKV E which consisted of the ectodomain of the ZIKV E protein fused with the transmembrane and cytoplasmic domains of the VSV-G protein (EG) (Fig 1). Pelleted supernatants from all conditions contained particles similar in size and morphology with typical MLV Gag particles as shown by ns-TEM analysis (Fig 2A) and both prME and EG plasmids induced expression of ZIKV E (Fig 2B). Preparations of EG-eVLPs contained a lower density of particles as observed in nsTEM (Fig 2A) but contained higher expression of the E protein than in prME eVLPs as indicated by Western blot analysis using anti-flavivirus mAb 4G2 (Fig 2B).

**Fig 1.**
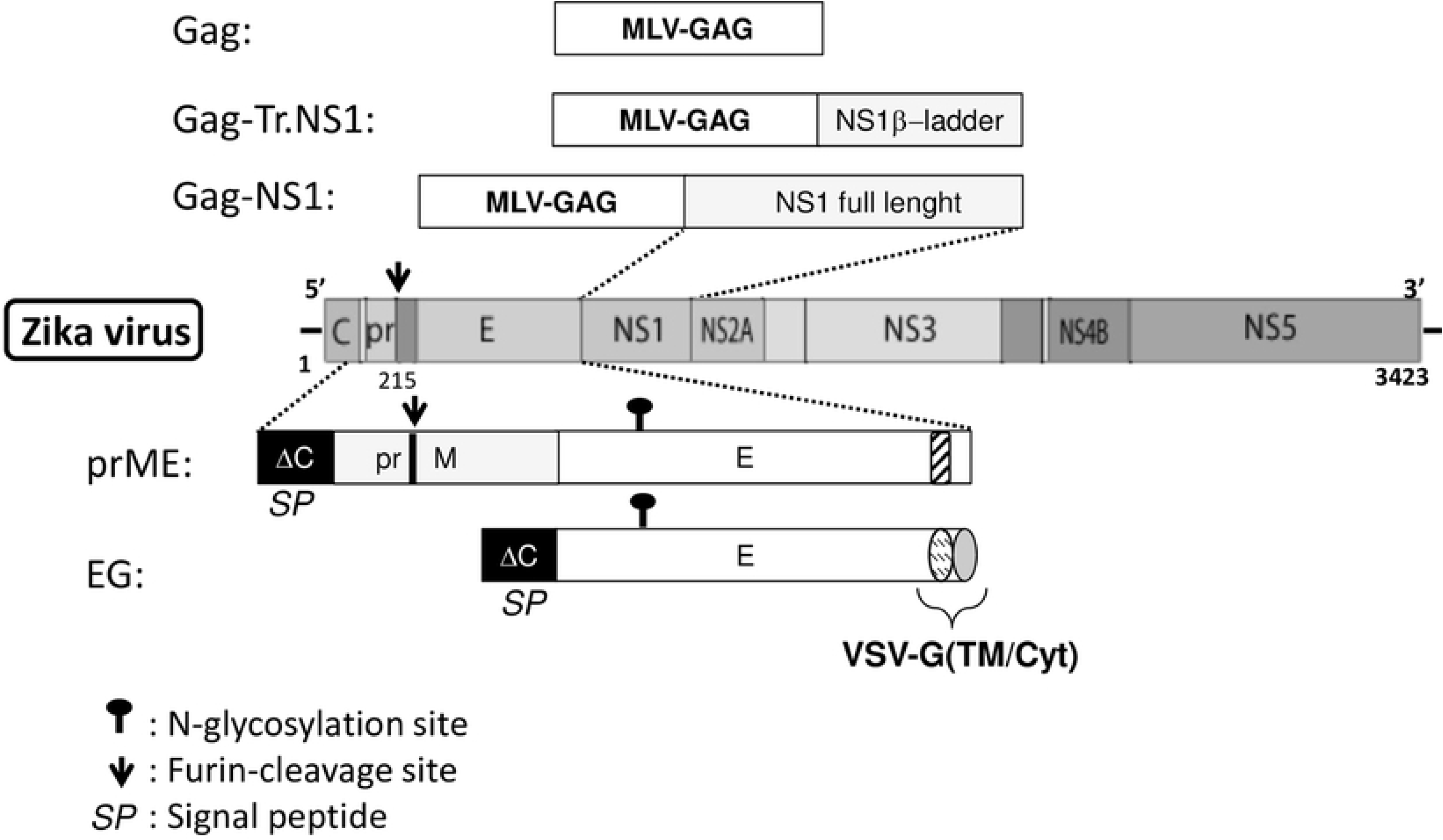
Schematic representation of the ZIKV genome and ZIKV plasmid constructs. prME design uses the unmodified full length sequence of E with 32 last aa of the capsid (ΔC) used as signal peptide (SP). In EG construct, the ectodomain of E (aa 291-744) has been fused with the transmembrane and cytoplasmic (TM/Cyt) domain of the VSV G protein (aa 468 to 511). The Gag sequence from Moloney murine leukemia virus (MLV-Gag) has been fused with either the full length non-structural NS1 (Gag-NS1) or a truncated NS1 (Gag-Tr.NS1) consisting of the β-ladder domain of NS1 (aa 180 to 353).

**Fig 2.**
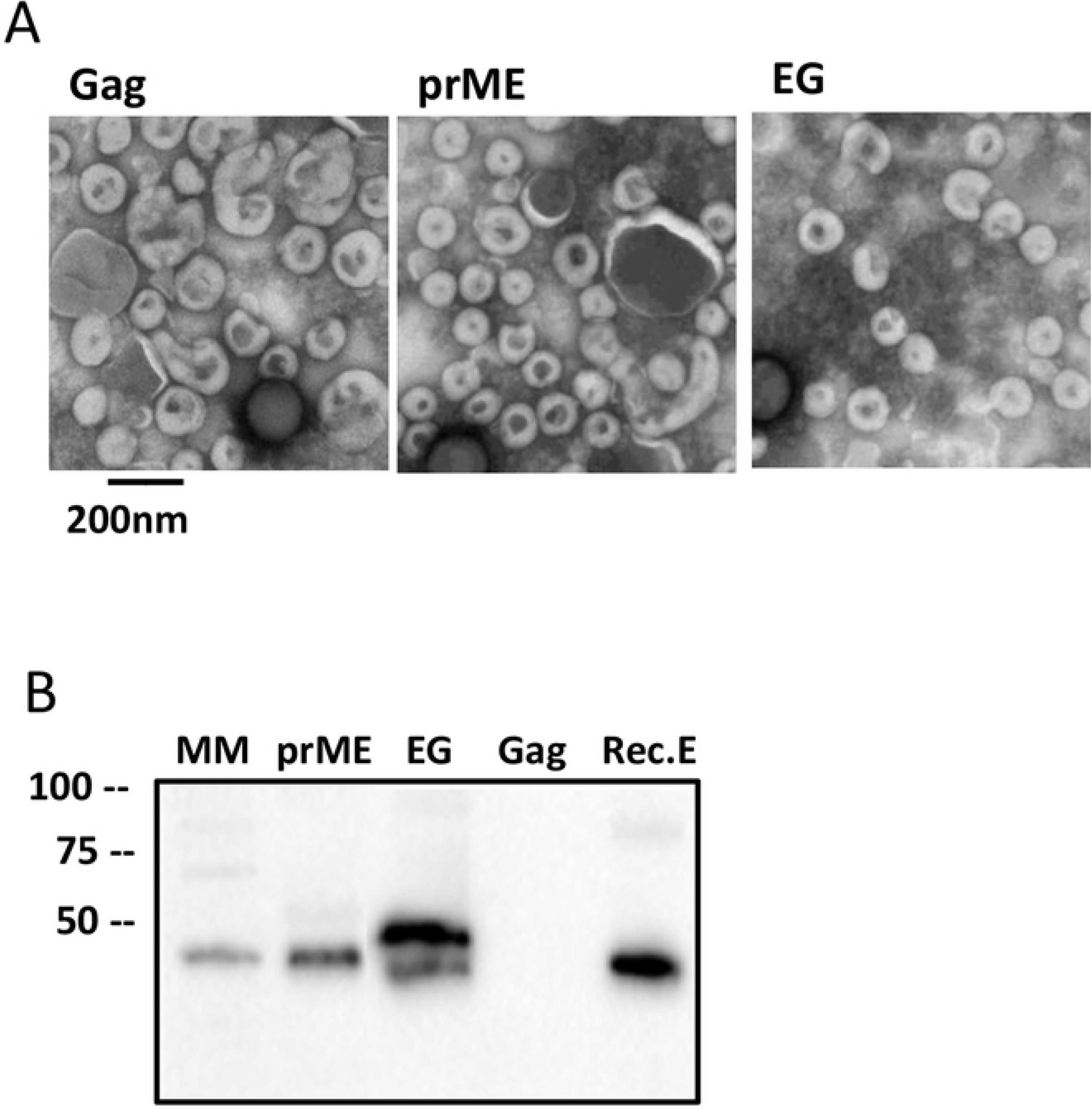
Characterization of ZIKV monovalent eVLPs expressing the E protein. **(A)** Representative nsTEM images at magnification x40,000x from eVLPs purified from culture supernatants of 293SF-3F6 cells transfected with Gag alone (Gag) or with Gag and ZIKV prME or EG constructs. **(B)** Expression of ZIKV E analyzed by Western-blot of ZIKV eVLPs and control using the anti-flavivirus mAb 4G2. eVLPs produced with Gag plasmid only (Gag eVLP) and recombinant ZIKV E (rec E) protein were used as negative and positive controls respectively.

When injected into BALB/C mice, EG eVLPs induced anti-ZIKV E IgG titers that were 100 to 1000 times greater than eVLPs expressing wild-type full length prME (Fig 3A) and in 7 of 8 mice, this strong IgG response was associated with a modest but significant neutralizing activity (Fig 3B). Based on this data, and reproducible yields of eVLPs with elevated levels of E protein contents, we selected EG as our preferred ZIKV E construct.

**Fig 3.**
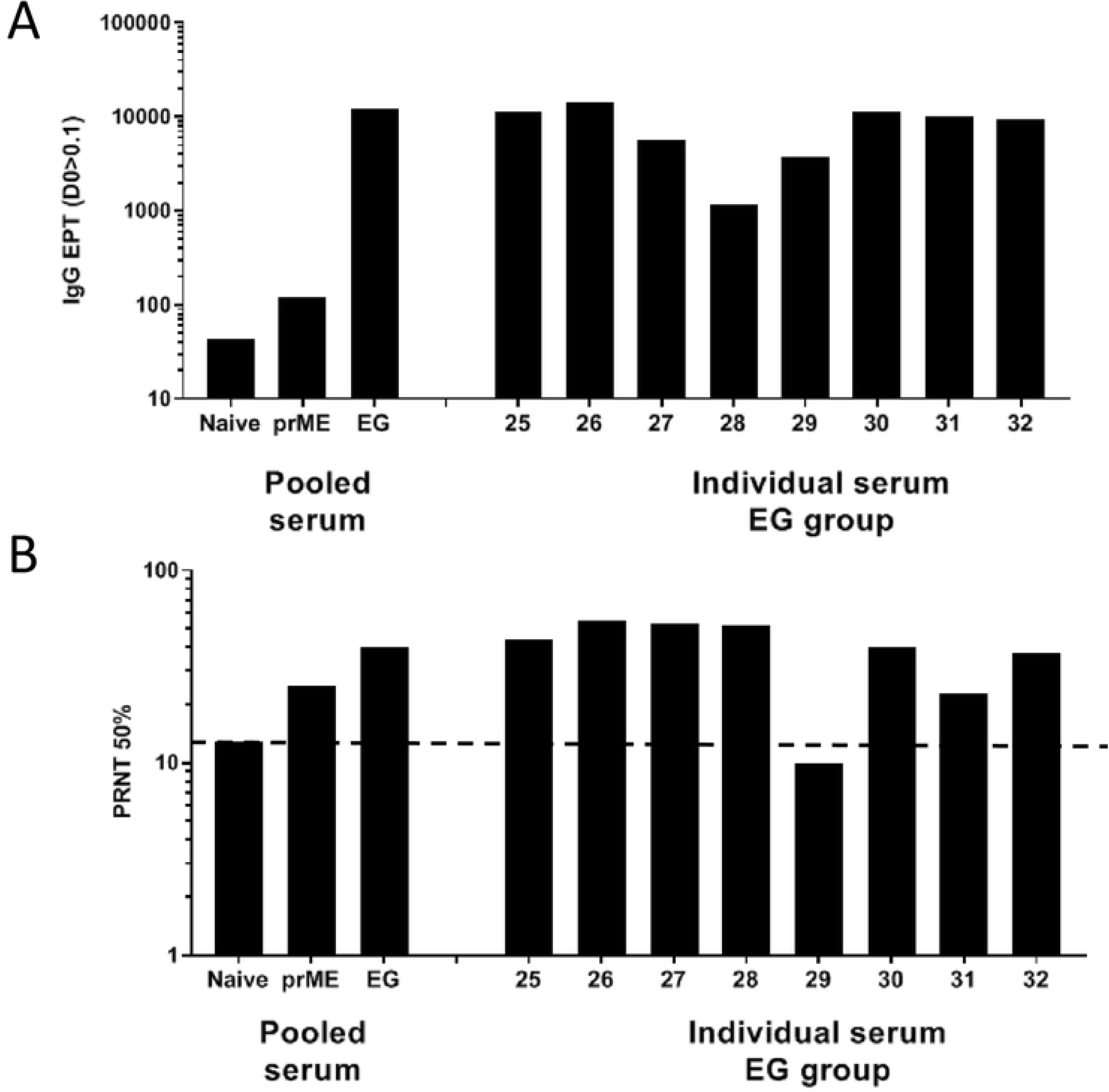
Immunogenicity of ZIKV E eVLPs. Four groups of 8 BALB/c mice received 2 injections of eVLPs at day 0 and 28. Sera were collected 4 weeks after the second injection and analyzed individually or as pooled sera. Sera pooled from 3 non immunized mice (naïve) were used to indicate background. **(A)** IgG end point titers (EPT) were evaluated in ELISA using recombinant ZIKV E protein. EPT was determined as the first dilution that gave an OD > 0.1. **(B)** The neutralization activity was determined in PRNT with a 50% threshold as described in Materials and Methods.

### Production and characterization of bivalent EG/NS1 eVLPs

In an attempt to potentiate the antibody neutralizing response to ZIKV via enhanced T cell help, we sought to develop bivalent eVLPs expressing NS1 protein as well as EG. As NS1 is readily secreted by infected cells, it was fused in-frame with Gag to ensure its retention inside the particles. The first set of experiment demonstrated that the full length NS1 fused to the full length Gag (Gag-NS1) resulted in very poor yields of eVLP particles, as indicated by very low levels of Gag protein detected by ELISA (Fig 4A). It is possible that the elongated size of the Gag-NS1 fusion protein, together with the specific properties of NS1, disrupted the architecture of the particles. This observation was consistent with our unpublished previous studies showing that eVLPs stability was more difficult to achieve when Gag was fused with an additional protein. A truncated form of NS1 corresponding to the hydrophobic β-ladder domain was chosen because its homologous domain in γ NS1 contains multiple CD8^+^ and CD4^+^ T cell epitopes (26,27). Transfection with Gag-Truncated NS1 (Gag-Tr.NS1) DNA plasmid produced particles with expected yields (Fig 4A). Co-transfection with Gag-Tr.NS1 and EG plasmids produced bivalent EG/NS1 eVLPs that expressed both and EG and Gag-fused NS1 protein as shown by western blot analysis (Fig 4B). Optimization of the amounts of plasmid DNA enabled to obtain similar amounts of Gag and particle numbers in bivalent and monovalent eVLPs for fair comparison of both preparations (Fig 4A). A standard procedure using 0.4 μg/mL of either Gag or Gag-Tr.NS1 plasmid was used for further studies.

**Fig 4.**
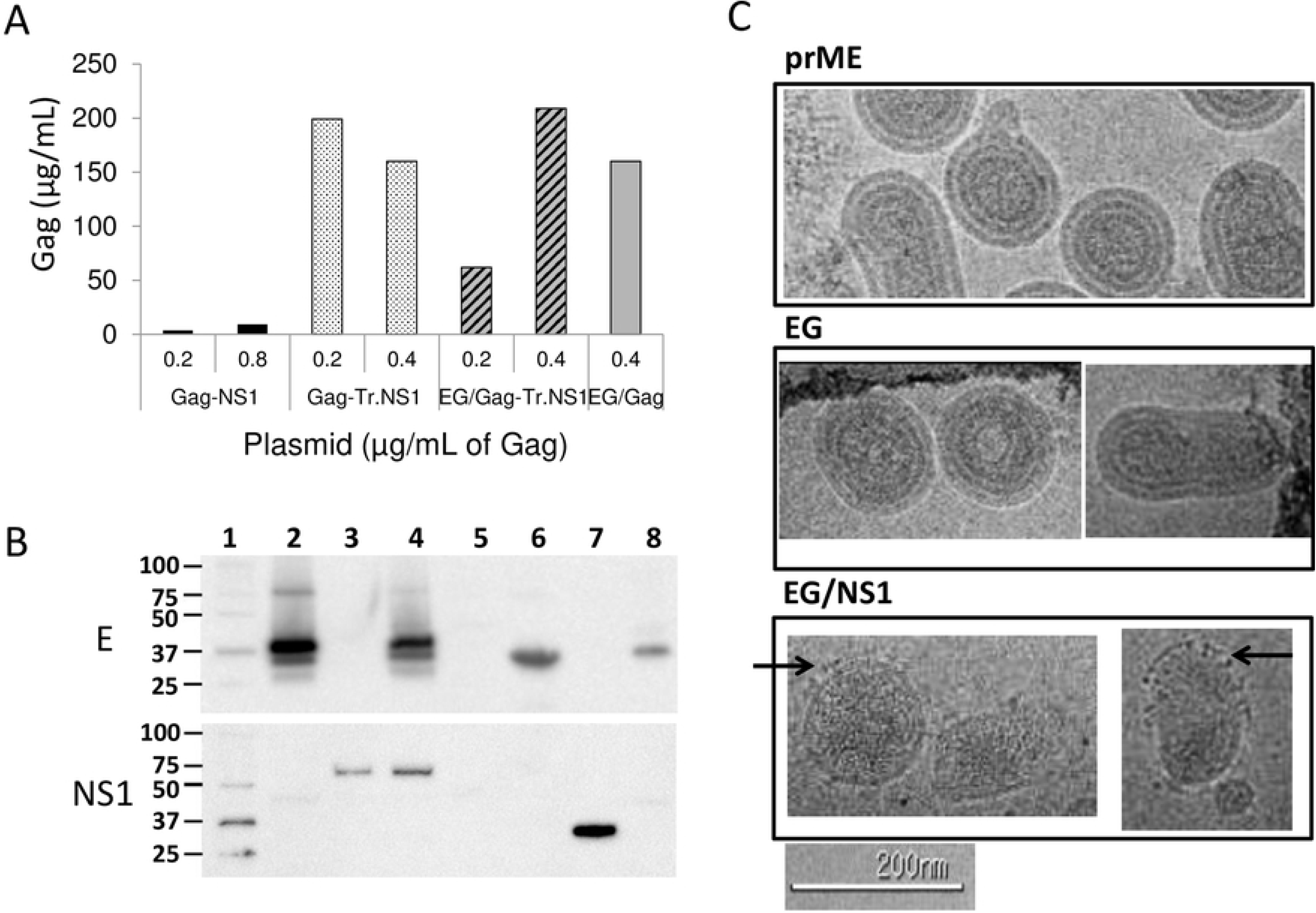
Production of bivalent eVLP expressing ZIKV E and NS1 proteins. **(A)** Gag amount was measured by SDS-PAGE densitometry in eVLPs preparations produced with 0.2 or 0.4 μg/mL of either Gag, Gag-NS1 or Gag-Tr.NS1 used alone or in combination with EG plasmid. Results are shown as the concentration of Gag in μg/mL in each preparation. **(B)** Western blot analysis was performed to detect E and NS1 protein in eVLPs preparations. 5μg of each test particles were loaded on acrylamide gel. Recombinant proteins are used for positive controls (rec.E; rec.NS1), and Gag eVLP produced with plasmid Gag alone were used as negative control. Western blot analysis was performed using anti-ZIKV NS1 mAb, and anti-flavivirus antigen mAb 4G2.1; 1, molecular weight ladder; 2, EG eVLPs; 3, NS1 eVLPS; 4, EG/NS1 eVLPs; 5, Gag eVLPs; 6, rec.E; 7, rec.NS1; 8, prME eVLPs. **(C)** CryoEM images from eVLPs expressing ZIKV proteins prME, EG or EG/NS1. EM imaging was performed from highly purified eVLPs samples using an FEI Tecnai T12 electron microscope, operating at 120keV equipped with an FEI Eagle 4k × 4k CCD camera. Vitreous ice grids were transferred into the electron microscope using a cryostage that maintains the grids at a temperature below −170 °C. Selected images at magnification of 21,000x (0.50 nm/pixel) are shown with scale bar 200nm. Arrows identify glycoprotein spikes on the surface of bivalent EG/NS1 eVLPs.

Monovalent EG and prME eVLPs had a different appearance on CryoEM images from bivalent EG/NS1 eVLPs (Fig 4C). Monovalent EG and prME eVLPs were multi-lamellar particles with smooth surfaces, comparable to the appearance of typical MLV Gag particles (28), with a smooth surface, like mature Zika virions (14). In contrast, bivalent EG/NS1 particles were predominantly uni-lamellar with a homogenous dense textured core and their outer surface contained structures with dark densities protruding ~8-10 nm from the lipid bilayers, and resembling spikes (Fig 4C). A similar impact on the core structure when using a plasmid in which MLV Gag was fused with the CMV pp65 protein has been observed (unpublished data). It is possible that the addition of a protein in frame with the Gag sequence induced an alteration of the architecture of the MLV-Gag VLP core structure (28), which may have prompted a conformation change in the surface EG protein due to altered interactions between the Gag/NS1 core and the cytoplasmic tail of the EG protein (29).

### Expression of NS1 β-ladder domain enhances the neutralizing activity of ZV EG eVLPs and induces T cell activation

BALB/C mice received 3 injections of eVLPs at monthly intervals after which the effect of NS1 on the response to EG eVLPs was evaluated. Mice immunized with bivalent EG/NS1 eVLPs produced comparable levels (p=0.48) of anti-ZIKV E IgG relative to mice immunized with monovalent EG eVLPs, but their sera demonstrated a higher proportion of neutralizing activity (p=0.05), comparable to naturally acquired immunity (Figs 5A-B). Sera from all groups contained very low to undetectable anti-NS1 IgG or IgM (Fig 5C). These results suggested that the antibody responses against NS1 were unlikely to contribute to the enhancement of neutralizing activity observed with the bivalent EG/NS1 eVLPs.

**Fig 5.**
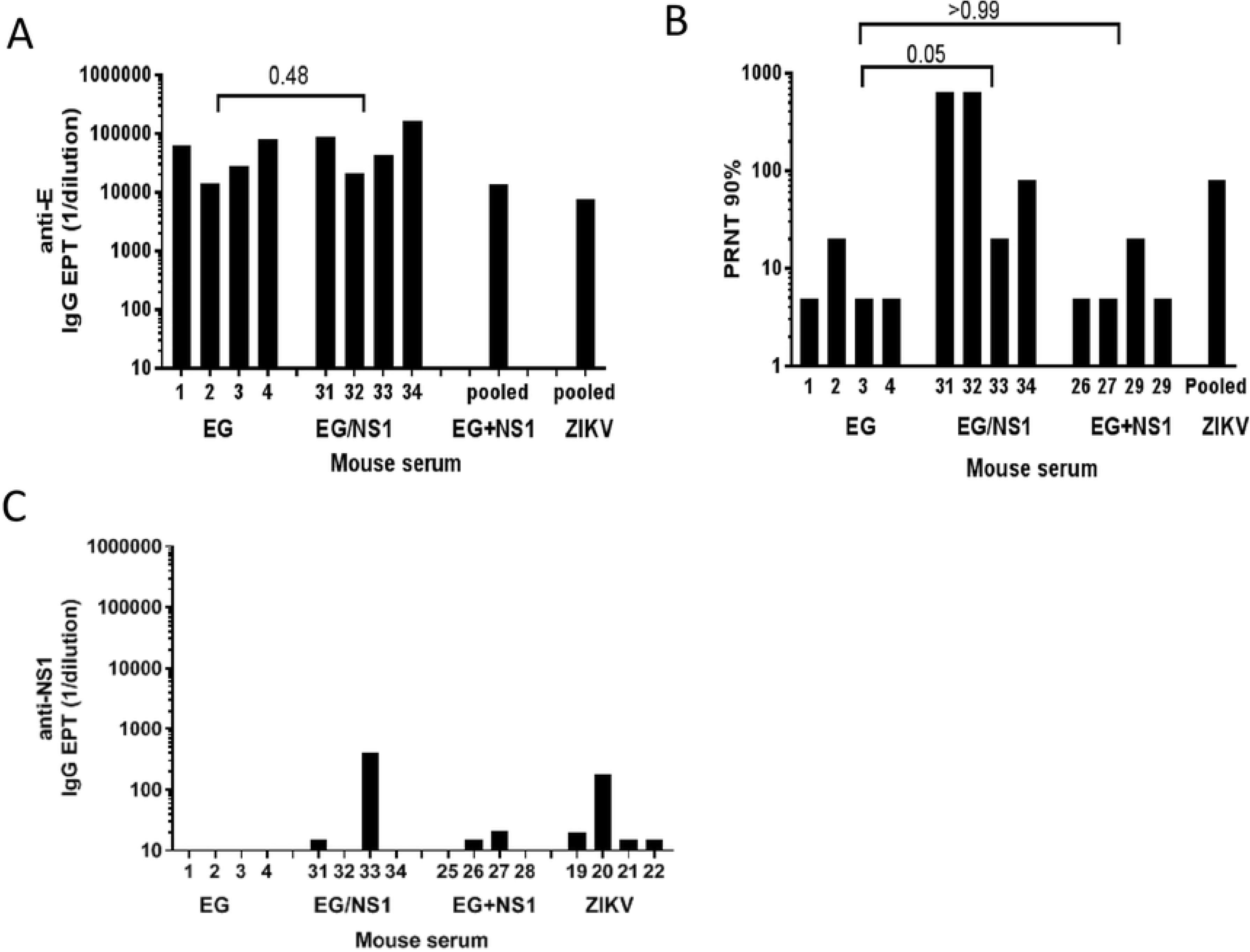
Antibody binding and neutralizing activity of anti-ZIKV serum. Three groups of 8 BALB/C mice received injections of eVLPs at day 0, 28 and 56. Groups were defined as follow: Group 1= EG eVLPs (EG), Group 2= combination of monovalent EG eVLPs + NS1 eVLPs (EG+NS1), Group 3= bivalent EG/NS1 eVLPs (EG/NS1). Amount of particles for each injection were adjusted to give 5μg of EG in all groups and 9μg of NS1 in groups 2 and 3. Anti-ZIKV antibody titers and neutralizing activity were measured at the end of the study (6 days after third injection). **(A)** anti-ZIKV E IgG EPT in individual sera from group EG and EG/NS1 and in pooled sera from EG+NS1 group and 3 mice recovering from ZIKV infection (ZIKV). **(B)** 50% plaque reduction neutralization titers (PRNT50%) in Vero cells were analyzed in the same samples than in (A). **(C)** anti-NS1 IgG EPT measured by ELISA against recombinant NS1 in individual sera. Statistical analysis was performed using non-parametric T test.

ZIKV NS1-specific T cell immunity, as shown by induction of IFN-γ and IL-5 T cell secretion (Figs 6A-B), was detected in all groups of mice that received NS1 eVLPs. Similar responses were observed when comparing monovalent NS1 eVLPs and bivalent EG/NS1 eVLPs. In contrast, co-injection of EG eVLPs with NS1 eVLPs did not enhance the nAb response and induced lower levels of T cell activation levels than those observed with EG/NS1. These results emphasized the value of having the EG and NS1 proteins co-expressed within the same particles.

**Fig 6.**
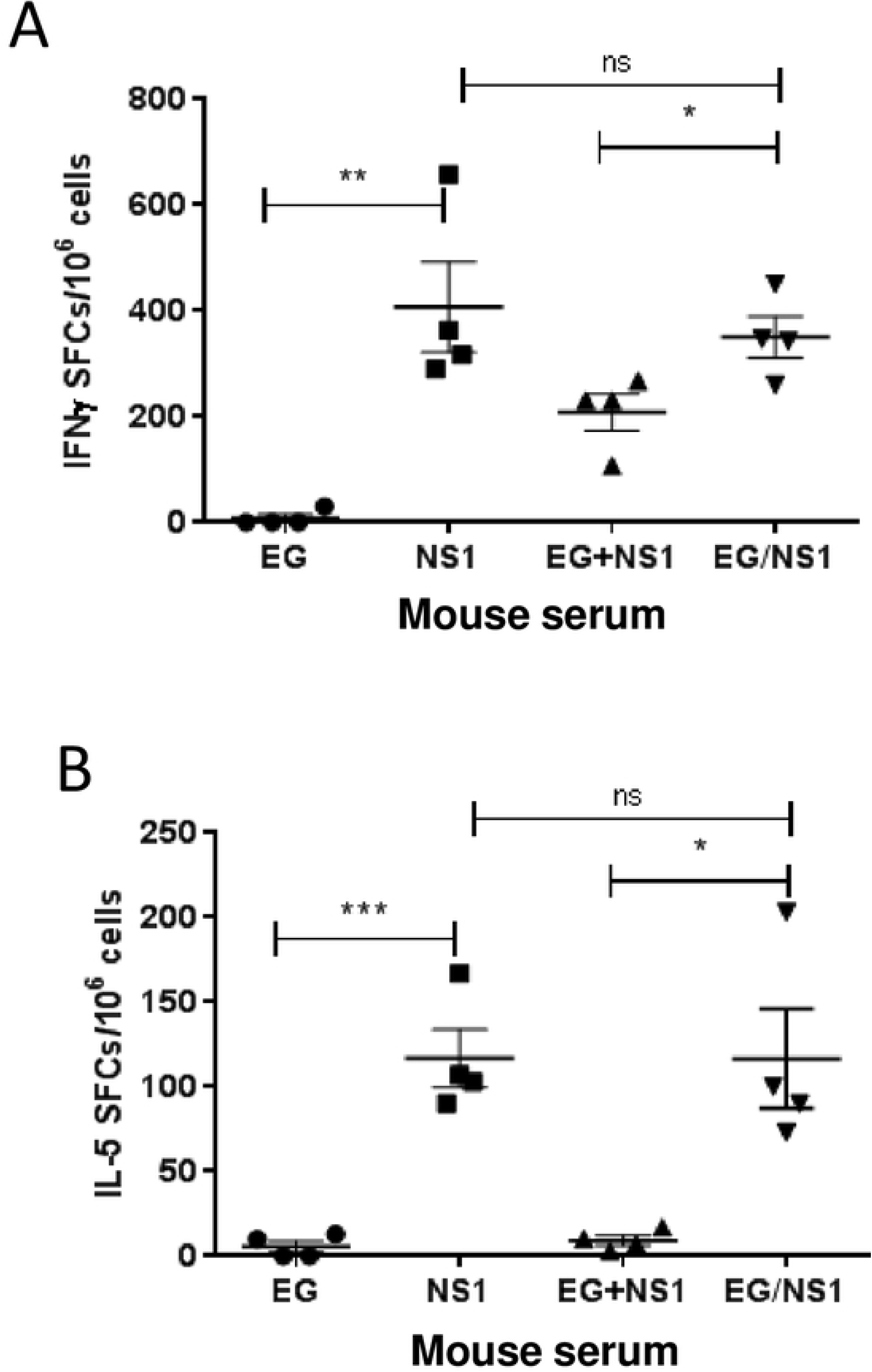
IFNγ and IL-5 production in BALB/C mice immunized with ZIKV eVLPs. At week 10, 4 mice per group randomly picked from the experiment described in Fig.5 were sacrificed and splenic T cells were cultured with NS1 peptides to measure the production of IFNγ **(A)** and IL-5 **(B)** by ELISPOT assay. Statistical analysis was performed using non-parametric T test.

### EG and EG/NS1 eVLPs protect mice from ZIKV challenge

The protective efficacy of the ZIKV eVLPs was examined in IFNαR deficient mice fully bred to a C57BL/6 background (C57BL/6-IFNαR^−/−^ mice). These mice, like BALB/C mice are able to mount protective immunity to ZIKV, but unlike the latter during primary infection, they also display significant weight loss associated with other overt sign of infection such as hind leg paralysis, the absence of which provides a facile read out of vaccine efficacy (30). Two separated challenge experiments were conducted successively using identical protocols with different groups.

A first challenge experiment was conducted to establish whether EG and EG/NS1 eVLPs could induce protective Ab immune response against ZIKV infection in the C57BL/6-IFNαR^−/−^ mouse model. Four groups of 10 mice were injected with either saline, eVLPs control expressing no ZIKV protein (Gag eVLP), EG eVLPs or EG/NS1 eVLPs. After a single injection of EG or EG/NS1 eVLPs, the levels of anti-E IgG rapidly increased in the serum of immunized mice peaking at 14 days after the second injection while the third injection maintained Ab titers at a plateau (Fig 7A).

**Fig 7.**
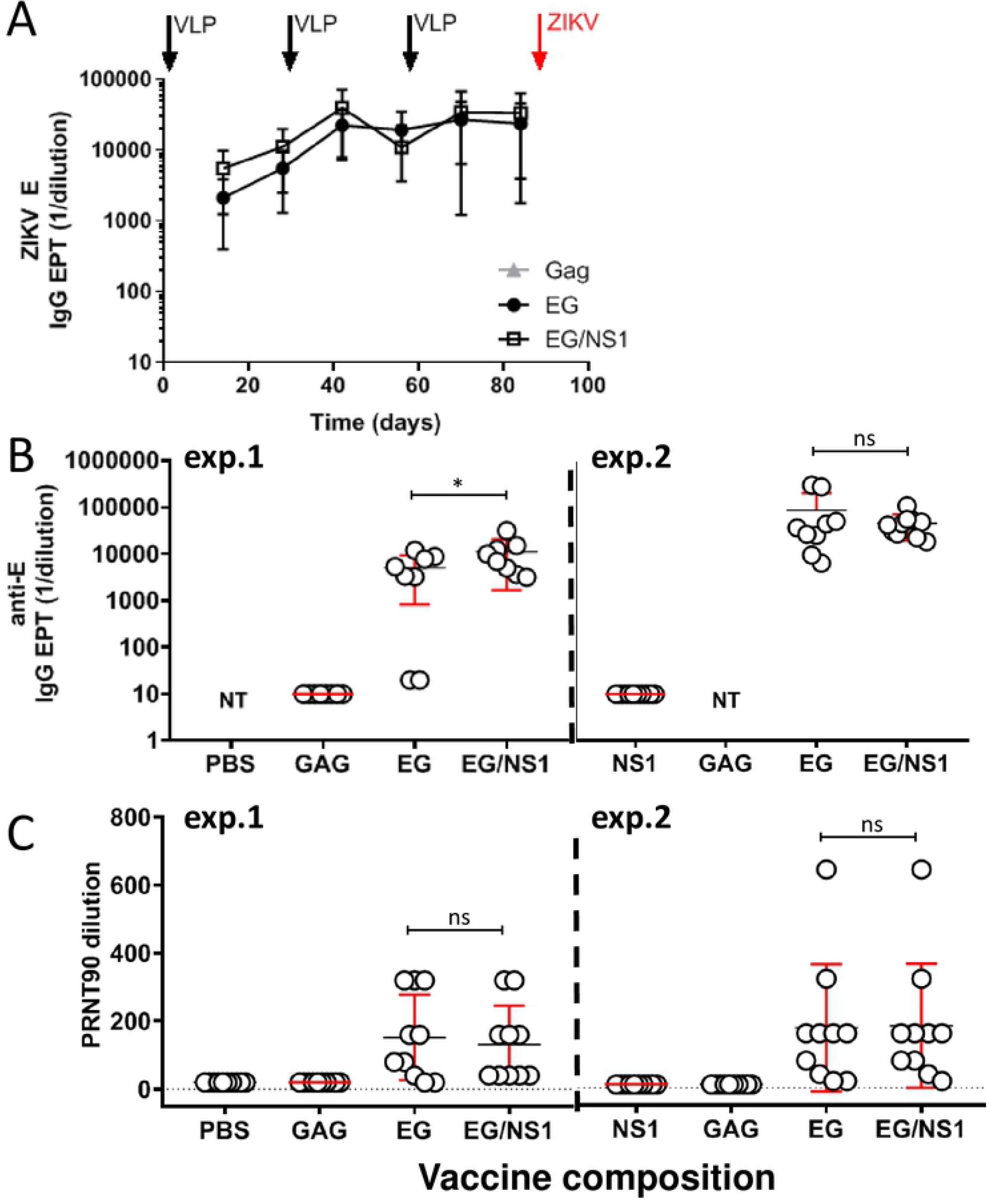
Induction of neutralization antibody response in C57BL/6αR^−/−^ mice after immunization with ZIKV eVLPs. Two experiments, exp.1 and exp.2, were conducted using a similar protocol. Four groups of 10 females C57BL/6αR1^−/−^ mice received injections of ZIKV eVLPs and control (saline or Gag eVLP) at day 0, 28 and 56. Sera were collected every 2 weeks. **(A)** Kinetics of anti-ZIKV E IgG titers measured by ELISA in individual sera from mice in exp.1. Data are expressed as mean +/− standard deviation of IgG EPT as a function of time. **(B)** Representation of IgG EPT measured in individual sera in both experiments using specific ELISA against ZIKV E protein. **(C)** Representation of neutralizing activity in individual sera in both experiments as evaluated by PRNT90% in Vero cells.

Evaluation of individual sera demonstrated that antibody responses were induced in 8 of 10 mice immunized with EG eVLPs and in 10 of 10 mice immunized with EG/NS1 eVLPs in Exp.1 (Fig 7B). Neutralization activity against ZIKV was evaluated by PRNT90% 27 days after the third injection. The sera from mice immunized with EG and EG/NS1 eVLPs possessed significant neutralizing activity which correlated with induction of antibody responses in the mice (Figs 7B-C). Of interest, the sera from both mice with undetected levels of anti-E had no neutralizing activity. A similar pattern of antibody response with higher levels of IgG was observed in a second experiment were a group of mice were injected with NS1 eVLPs instead of saline. In this experiment, all mice EG and EG/NS1 groups develop an Ab response. We further analyzed the immunoglobin isotype usage in the sera from mice from Exp.1. The IgG1 to IgG2b ratio indicated a predominant Th2 response in the EG group as expected from Alum adjuvant. However a significant difference was observed with bivalent EG/NS1 eVLPs, which promoted a switch towards IgG2b, that is usually associated with cell-mediated immunity (Fig 8).

**Fig 8.**
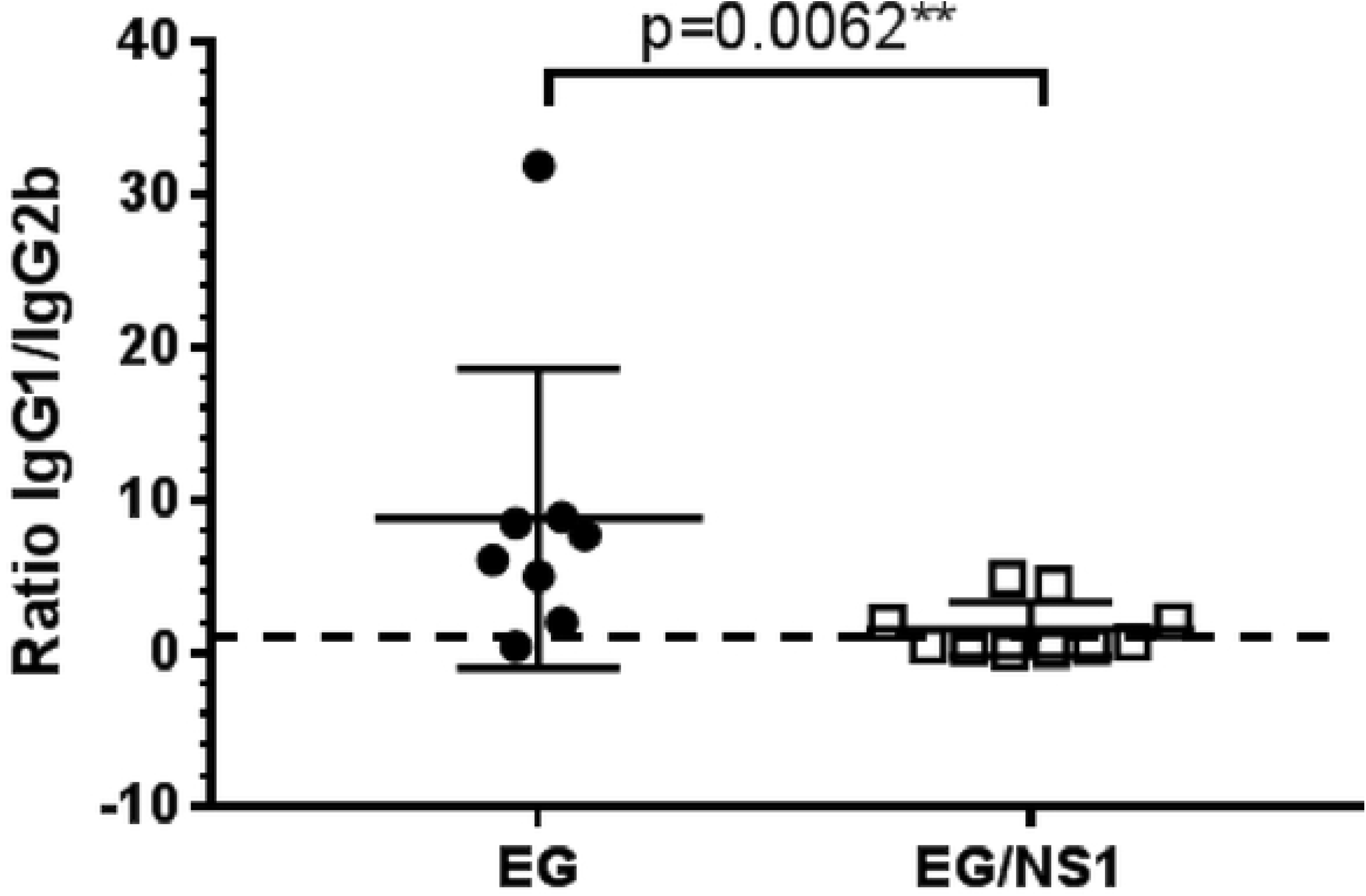
Ratio of IgG1/IgG2b anti-ZIKV E in mice immunized with ZIKV eVLPs. Isotype usage was determined by specific ELISA using HRP conjugate goat Ab against mouse IgG1 and IgG2 (Bethyl). Results are expressed as the ratio of IgG1 to IgG2b. Statistical analysis was performed using a non-parametric test.

After ZIKV challenge, mice were monitored daily for body weight change and behavior. C57BL/6-IFNαR^−/−^ mice in groups that received saline, Gag or NS1 eVLPs lost body weight and developed hind leg paresis between days 8-10 after ZIKV injection (Fig 9). In marked contrast, in both experiments, mice immunized with either monovalent EG or bivalent EG/NS1 ZIKV eVLPs showed only transient weight loss up to day 5 (Fig.9A-B), at which point they began to regain normal body weight (achieved at day 10). Although both monovalent EG eVLPs and bivalent EG/NS1 eVLPs groups showed statistically comparable levels of protection and total IgG, 2/10 mice in the EG eVLPs group in exp.1 and 1/10 in exp.2 (Fig 9C), developed hind leg paresis while none of the mice in the EG/NS1 group in both experiments developed paresis at any time. Of interest, no significant anti-ZIKV IgG Ab titer nor neutralizing activity was detected in neither of both non-protected mice from exp.1.

**Fig 9.**
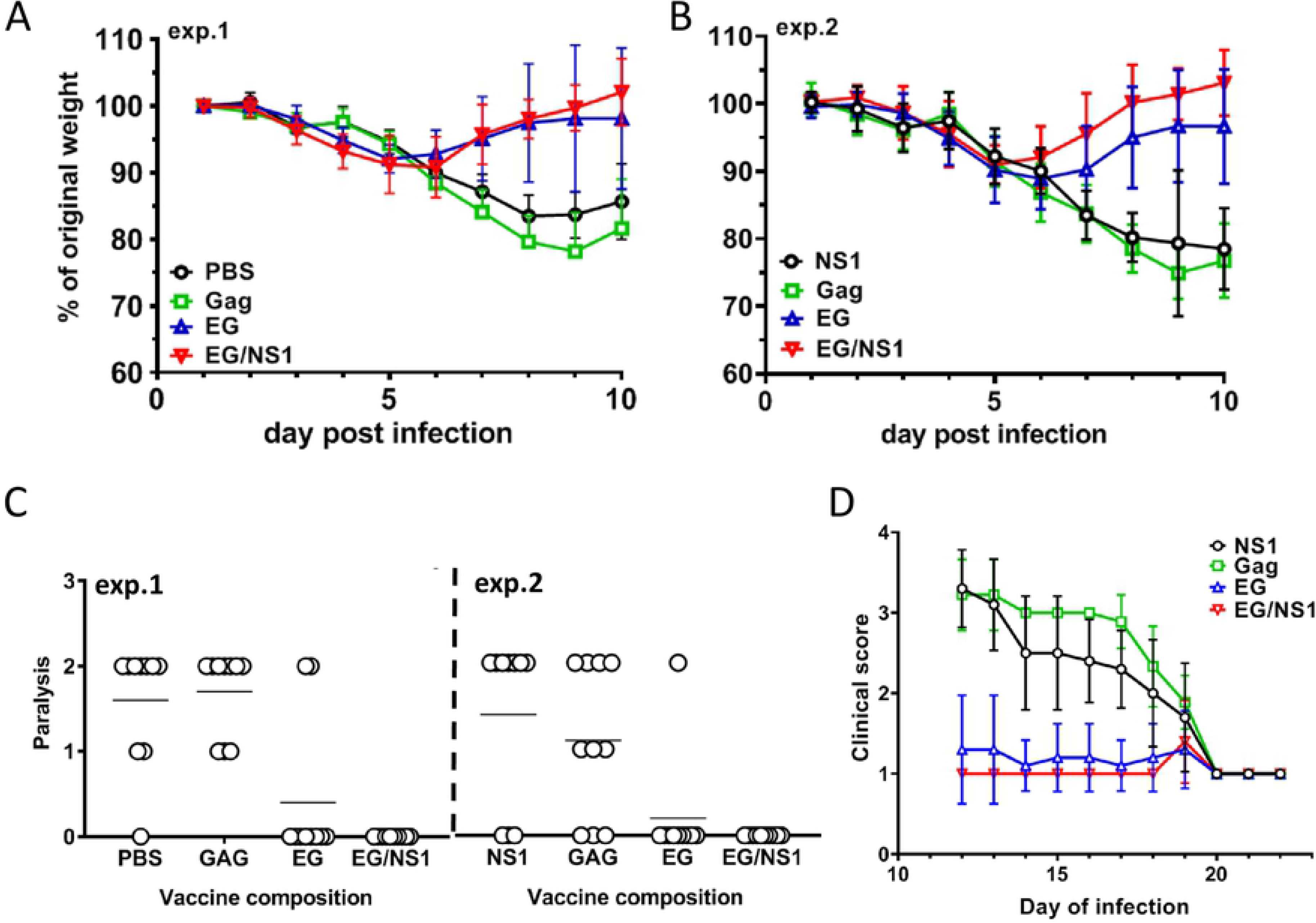
Clinical evaluation of immunized mice after ZIKV injection. Mice from exp.1 and exp.2 described on Fig 7 were inoculated with 10,000 PFU of ZIKV Philippine 2013 at 4 weeks after final injections or eVLPs and control PBS. **(A-B)** Mice were monitored daily for weight change. The figure represents the kinetic of weight change from ZIKV injection to day 10 after injection. **(C)** Each point on the figure represent the highest degree of paralysis scored for each mouse during the full course of daily observation from day 8 to day 11 post challenge. Hind leg paralysis was scored as follow: 0= no paralysis, 1= unilateral paralysis, 2= bilateral paralysis. Statistics were performed using Kruskal-Wallis non-parametric test followed by Dunn’ multiple comparisons test. Adjusted p values are shown.

The second challenge experiment was aimed at determining if NS1 eVLPs used alone could induce protection despite the absence of an efficient Ab response. Careful observation of animal behavior showed that despite no significant protection from paralysis compared to the control group (Fig 9C), some mice injected with NS1 eVLPs had milder symptoms after day 10 (Fig 9D).

## Discussion

Several ZIKV vaccine candidates have advanced through efficacy studies in mice and non-human primates (NHP) and are either in, or poised for, Phase I clinical trials. These include DNA-based vaccines expressing the ZIKV envelope protein (EP), whole killed ZIKV, and recombinant viruses including recombinant DENV and YFV expressing ZIKV EP (31–34). Live-attenuated and replication-competent vaccines are expected to provide long-term immunity and to be highly effective (35), but these vaccines are usually contraindicated in immunocompromised individuals and in pregnant women because of a potential risk of reversion of pathogenicity. The development of Zika vaccines may also be complicated by potential issues of ADE mostly due to a high degree of homology among the members of this family, especially between ZIKV and DENV (36).

We have developed an eVLP platform that led to the production of a CMV vaccine candidate currently under Phase I clinical trial evaluation (37). In the present study we have used this eVLP platform to develop a prophylactic ZIKV vaccine candidate that includes a modified form of the ZIKV E protein, consisting of swapping its transmembrane/cytoplasmic domain for that of the VSV-G protein as well as the β-ladder domain of the NS1 protein. The present study demonstrated that co-expression of the β-ladder domain of NS1 provides a synergistic effect on the neutralizing antibody response to the ZIKV envelop protein, and is associated with significant T cell activation

In immunocompetent BALB/C mice, EG eVLPs injected in the presence of Alum adjuvant induced a potent IgG response but neutralizing activity was modest. We hypothesized that NS1 could induce NS1-specific T-cell activation to provide T-cell help and increased neutralizing activity. When NS1 and EG were co-expressed in EG/NS1 eVLPs, a synergistic effect was observed with the neutralizing IgG response to reach an average of 100 fold increase over the response to EG alone. Mice that received eVLPs expressing NS1 showed a marked increase in NS1 specific T-cell activation that correlated with the increase in anti-ZIKV E IgG (r=0.9).

Despite the presence of several B-cell epitopes in the sequence used in our NS1 construct (38,39), NS1 and EG/NS1 eVLPs induced no or low Ab responses to this protein. In contrast, previous studies have demonstrated that vaccine candidates expressing full length NS1 promoted efficient specific anti-NS1 neutralizing Ab responses associated with protection against ZIKV infection (12,40–42). The discrepancy between our data and others is likely due to alteration of NS1 presentation in two ways: i) the NS1 domain was fused with Gag which can alter the tri-dimensional structure of NS1, ii) the NS1 domain is kept inside the particles, reducing its accessibility to B cells. These data reveal another potential utility of eVLPs for the analysis of T-cell versus B-cell responses against proteins, depending on their location on or within the particles. Of note, the β-ladder of the NS1 is flexible and can interact with many proteins. It is then possible that the use of a fusion protein Gag-NS1 can induce alteration of the structure of the particle, and subsequent alteration of the coexpressed EG protein, as suggested by the spike-like appearance of particles analyzed by cryo-EM (29).

As observed in BALB/C mice, EG and EG/NS1 eVLPs induced similar levels of anti-E IgG in C57BL/6-IFNαR^−/−^ mouse sera. However, in C57BL/6-IFNαR^−/−^ mice, expression of EG alone was sufficient to induce a strong neutralization antibody response equivalent to the one observed in response to EG/NS1 eVLPs in 18 out of 20 mice across 2 independent studies. Future studies may clarify if this difference is due to the genetic differences between C57/BL6 and BALB/6, or to the IFNαR pathway deficiency in C57/BL6-IFNαR^−/−^. All mice that mounted a nAb response, either in response to EG eVLPs or EG/NS1 eVLPs were protected against experimental ZIKV infection.

These data confirm that neutralizing Ab responses are critical in protection against ZIKV clinical disease and that the bivalent EG/NS1 eVLPs provide a more robust induction of nAb response than monovalent EG eVLPs. In C57BL/6-IFNαR^−/−^ mice, NS1 promoted a switch towards a Th1 antibody response, emphasizing a strong effect on T-cell help even in the presence of the Th2-adjuvant Alum.

It has been shown that an Adenovector ZIKV vaccine that induced a T-cell response against the ZIKV envelop without any Ab response could protect C57BL/6 mice against ZIKV infection (43). In our model, the C57BL/6-IFNαR^−/−^ mice that received NS1 eVLP alone were not protected from ZIKV infection in absence of an efficient nAb response. Further studies would be necessary to determine the role of T cells in these IFNαR deficient mice.

Flaviviruses possess a high degree of homology that leads to the abundant production of cross-reactive Abs. While the mechanism for ADE has not been fully elucidated, it is generally recognized that the presence of cross-reactive Abs with low neutralizing potency are involved in ADE (15). The neutralization potency depends on the affinity of the Abs and on the accessibility of the epitopes on the virions (44,45). In the present study, mice immunized developed a high neutralizing response that correlated with protection. Most potent nAb with low/no ADE have been described to be directed against EDIII domain and quaternary epitopes from the DI/DII domain (46,47). Further structural studies would determine whether the EG conformation at the surface of the eVLPs, would present such epitopes in an optimal conformation for Ab production.

VLPs expressing vaccine antigens are typically more immunogenic than monomeric recombinant forms of vaccine antigens because of repeating, array-like presentation of antigens, which is a preferred means of activating a B cell response. Moreover, the particulate nature of the antigen, relative to recombinant proteins is a much better means of activating dendritic cell responses and further enhances immunity. While being a highly immunogenic means of delivering vaccine antigens, VLP-based vaccines avoid the potential safety concerns associated with whole killed or recombinant viral-based vaccines, as there is no residual host DNA/RNA and no possibility for infection.

When glycoproteins are expressed/present on the surface of eVLPs, one terminus of the protein is anchored within the lipid bilayer, which imposes structural constraints not readily observed with monomeric recombinant antigens. Moreover, by altering the transmembrane domain/cytoplasmic tail of the glycoprotein, and/or the core particle structure with which the cytoplasmic tail may interact, we have found that a novel presentation of glycoproteins can be obtained that is associated with enrichment for antibodies with neutralizing rather than just binding activity. This was demonstrated previously with eVLP expression of a modified form of the CMV gB (25), similar to what we have observed with the EG/NS1 ZIKV vaccine candidate presented in this study. The high degree of neutralization obtained with EG/NS1 eVLPs may minimize the risk of ADE and is an attractive candidate for further development as a prophylactic vaccine against ZIKV.

## Material & Methods

### Cells and viruses

293SF-3F6 cell line derived from human embryonic kidney (HEK) 293 cells is a proprietary suspension cell culture provided by the National Research Council (NRC, Montreal, Canada) and grown in serum-free chemically defined media (22). All other cell types (Vero) and viruses (ZIKV strain PRVABC59 and Philippines 2013) were grown and handled by NRC or Southern Research Institute according to their internal SOPs based on American Type Culture Collection recommendations.

### Construction of plasmids

We used the ZIKV E sequence from Suriname isolate 2015 (23) (Genbank KU312312) to construct a plasmid expressing the full length unmodified ZIKV prME and a plasmid expressing the ectodomain of E fused with the transmembrane and cytoplasmic domain of the VSV G protein (EG) (24). The ZIKV E sequences were preceded by a portion of the sequence corresponding to the last 32 aa from the Zika virus capsid, as signal peptide. For the production of VLP, we used a minimal cDNA sequence encoding a Gag polyprotein of murine leukemia virus (MLV) (Gag without its C-terminal Pol sequence) or Gag-NS1 or Gag-TR.NS1, were Gag was fused with full length NS1 or truncated NS1 consisting of the β-ladder domain of NS1 (aa 180 to 353). The design of the plasmids is summarized in Fig1. All final sequences were synthetized and optimized for mammalian cell expression at Genscript (NJ), prior to subcloning into our in-house phCMV-expression plasmid (24).

All DNA plasmids were amplified in high-efficiency competent *Escherichia coli* cells and purified with an endotoxin-free preparation kit using standard methodologies.

### Production and purification of eVLPs

eVLPs were produced using transient polyethylenimine transfection in HEK 293SF-3F6 GMP compliant cells as described previously (25). Harvests containing eVLPs were purified either by ultracentrifugation using sucrose cushion as described previously (25) or by a proprietary method that consists of tangential flow filtration concentration, diafiltration and ultracentrifugation using sucrose cushion. The final product was sterile filtered using 0.2 μm membrane prior to formulation and injection into animals.

### Characterization and quantification of protein contents

MLV-Gag, NS1 and E protein content were analyzed by western blotting as described previously (25). using mouse monoclonal anti-flavivirus group antigen clone D1-4G2-4-15 (EMD Millipore) for detection of ZIKV E and mouse anti-Zika NS1 mAb (BioFront Technologies) as primary Abs, and goat anti-mouse HRP conjugate (Bethyl). Precision Protein Streptactin HRP conjugate (Bio-Rad) was used as molecular weight ladder and recombinant ZIKV E and NS1 protein as standard controls (Meridian Life Sciences). Total protein concentration in eVLP samples was determined using Bicinchoninic Acid (BCA) protein assay according to the manufacturer’s instructions (Thermo-fisher Scientific). SDS-PAGE separation followed by Coomassie staining was performed to determine the total amount of E, and NS1 proteins in the eVLP samples. Briefly, protein in the samples were denatured and reduced with commercial Laemmli buffer supplemented with 50 mM DTT in a boiling water bath. Proteins were then separated in a pre-cast 4-20% polyacrylamide gradient gel, stained with commercial Coomassie stain and then de-stained per the manufacturer’s instructions (BioRad). All image processing and densitometry was performed with ImageLab software (BioRad). The E and NS1 contents of the eVLP samples were quantified by densitometry analysis of E, and NS1 bands that were identified separately by western blots using monoclonal antibody (see section above). The banding patterns and the protein contents of the eVLP samples were determined using BSA standard (BioRad). The MLV Gag content in each eVLP preparations was quantified using a proprietary sandwich ELISA. Briefly, the assay was performed using a combination of monoclonal antibody R87 to MLV-p30 as capture, and a rabbit polyclonal anti-MLV p30 as detection antibody, with commercially recombinant Gag as standard antigen. The samples were treated for 20 min with 0.5% SDS followed by 0.5% Triton-X for 30 min at room temperature. HRP-conjugated anti-MLV p30 polyclonal antibody was used as detection antibody and the reaction was terminated by addition of stop solution and the absorbance was measured at 450 nm. The data fitting and analysis are performed with Softmax Pro 5, using a four-parameter fitting algorithm.

### Electron microscopy

Negative staining transmission electron microscopy (nsTEM) analysis of the eVLPs was performed at the Armand-Frappier Institute (INRS) in Montreal, Canada following standard procedures. Cryo-TEM was performed at NanoImaging Services in San Diego, CA according to standard procedures.

### Animal experiments

BALB/c mice were purchased from Charles River Laboratories (Saint-Constant, Quebec). The animals were allowed to acclimatize for a period of at least 7 days before any procedures were performed. IFNαR−/− mice have been backcrossed to a C57BL/6 background and bred at the National Research Council of Canada (NRC) animal care facility.

Four to six-week old female mice, 8 to 10 per group depending on experiments, received 3 intraperitoneal (IP) injections of eVLPs formulated in aluminum phosphate gel, Alum Adjuvant (Adju-Phos®, Brenntag) or saline control at monthly intervals or as indicated for each experiment. Blood samples were collected every 2 weeks starting 2 weeks before the first injection. Sera were either pooled or analyzed individually.

Challenge with 10,000 PFU of ZIKV Philippine 2013 was done 4 weeks after final injection in C57BL/6-IFNαR^−/−^ mice by the IP route. After infection, the mice were monitored daily for body weight and clinical behavior as indicated in figure legends and blood was collected at the end of the study.

### Monitoring of immune response

Direct ELISA described earlier (25) was used to measure Ab binding titers to recombinant E and NS1 protein from ZIKV (both from Meridian LifeScience). Plaque Reduction Neutralization Test (PRNT) was used to measure ZIKV-specific neutralizing activity in mouse sera. Vero cells were seeded at 3×10^5^ cells/mL in 24-well plates 24h prior PRNT. On the day of assay, virus and serially diluted serum samples were mixed and incubated for 1h at 37±1°C. Supernatant from cell-seeded 24-well plates was decanted then 100 ml of virus/serum mixture was transferred from the dilution plate to the cells. After 1h adsorption, agarose-containing overlay media was added and plates were incubated at 37°C, 5%CO2 for 3 days. The cells were fixed and stained using crystal violet solution and plaques were counted visually. The neutralizing antibody titer was expressed as the highest test serum dilution for which the virus infectivity is reduced by 50% or by 90%, as indicated in legends. NS1-specific IFNγ and IL-5 production by ZIKV-specific splenic T cells were enumerated by Enzyme Linked ImmunoSpot (ELISPOT) assay following manufacturer’s instructions (Mouse IFN-γ/IL-5 ELISpot Basic kits; Mabtech). Mouse splenocytes were left unstimulated or stimulated with an NS1 peptide pool (PepMix ZIKV NS1 Ultra; JPT Peptide Technologies, Germany) designed to cover the high sequence diversity of ZIKV. Spots were counted at ZellNet Consulting (NJ, USA) using KS ELISPOT Reader System (Carl Zeiss, Thornwood, NY, USA) with KZ ELISPOT Software Version 4.9.16. The background was defined as the number of spot forming cells (SFCs) per 10^6^ splenocytes in non-stimulated samples.

### Ethics Statement

All animal (mice) studies were conducted under ethics protocols (2010.17 and 2017.07), which was approved by the NRC Animal Care Committee. The mice were maintained in a controlled environment in accordance with the Canadian Council on animal care “Guide to the care and use of experimental animals” at the NRC Animal Research facility (Ottawa, Canada).

## Acknowledgements

We thank NRC animal resource staff (Human Health Therapeutics Research Centre) and Craig Bihun for assistance in animal studies.

We thank Diana Duque, Adam Asselin, Stephanie Pichette and Matthew York for their valuable contributions to the work described here.

This work is dedicated to the memory of Dr. Judie Alimonti (1962-2017) who began the ZIKV studies at NRC that led to the work reported herein.

